# Artificial hibernation uncovers distinct interaction dynamics in functional and pathological state change

**DOI:** 10.64898/2026.10.19.759973

**Authors:** Damien Mercier, Ming-Ching Chiang, Viviane M. Saito, Yu-Ju Lin, Hiroto Ashitomi, Minami Oshiro, Ai Takahashi-Nakazato, Kazumasa Z. Tanaka

## Abstract

Neural population activity can reorganize without disrupting function, but how such changes differ from those associated with pathology remains unknown. Artificial hibernation, Q-neuron-induced hypothermia and hypometabolism (QIH), causes extensive synaptic remodeling in the hippocampus without erasing established memories, allowing us to examine how extensive network reorganization can occur without lasting functional impairment. Longitudinal calcium imaging showed that hippocampal activity recovered after QIH without returning to its pre-hibernation population configuration. Instead, QIH reorganized neuronal activity patterns and coactivity beyond representational drift. Electrophysiological recordings revealed that this reorganization of neuronal interactions depended on the preceding network state. In mice subjected to kainic acid-induced status epilepticus, QIH reduced seizure-associated increases in neuronal interactions and spike coactivity during sharp-wave ripples. Over the following 50 days, the divergence of hippocampal LFP trajectories was constrained, and the progressive development of electrophysiological and histological pathology was attenuated. These findings suggest that controlled, extensive reorganization of neural population activity can prevent the progression toward pathological network states, with changes in neuronal interactions, particularly sharp-wave ripple-associated coactivity, accompanying this process.

## Introduction

Stable perception, memory and behavior have traditionally been assumed to depend on correspondingly stable patterns of neural activity. Longitudinal recordings have instead revealed that the neurons participating in a representation, their response properties and the resulting population activity patterns can change continuously over days to weeks, even under unchanged sensory conditions and without apparent disruption of memory or behavioral performance. This phenomenon, termed representational drift, has been observed in hippocampal spatial representations, association cortex and primary sensory cortices ^1–7^. Despite substantial changes at the level of individual neurons, task-relevant information and statistical features of population activity can remain preserved. Stable function therefore does not require the brain to reproduce a unique population configuration, but can be supported by neural representations that are continually reorganized over time.

Continuous reorganization of neural population activity, however, does not always preserve function. Progressive changes in population activity are also observed during the development of neurological disorders, including epilepsy and neurodegenerative diseases ^8–10^. These changes involve alterations in neuronal interactions, population activity patterns, and network dynamics, and can precede or accompany the emergence of functional impairment ^11^. For example, longitudinal recordings during epileptogenesis have revealed progressive changes in functional connectivity and network dynamics, even in background activity outside overt seizures ^8^. Thus, continuous reorganization of neural activity can occur both when function remains stable and when it deteriorates. However, what distinguishes the trajectories of population reorganization associated with these different functional outcomes remains poorly understood.

Artificial hibernation provides a model to examine how neural population activity evolves through an extreme physiological perturbation without lasting impairment of brain function. Under physiological conditions, prolonged suppression of neuronal activity can trigger homeostatic plasticity, including synaptic scaling that compensates for reduced activity ^12–14^. In contrast, Q-neuron-induced hypothermia and hypometabolism (QIH) produces profound suppression of hippocampal neuronal firing and network oscillations, including theta, gamma and sharp-wave ripples, lasting for up to 48 hours without compensatory recovery of neuronal activity ^15–17^. Despite this prolonged suppression and extensive synaptic remodeling, animals recover without overt behavioral abnormalities, memory impairment and deteriorated hippocampal spatial representations acquired prior to QIH. The preservation of brain function after QIH is consistent with the idea that neural population activity resumes its preceding functional trajectory. Characterizing population activity during QIH and subsequent recovery may therefore reveal the extent and nature of neural reorganization that can occur without compromising its function.

A critical question is whether the preservation of function after QIH also reflects preservation of the preceding trajectory of neural population activity. Even substantial, yet functional, changes in population configuration could reflect accelerated representational drift without altering the subsequent course of network evolution. Conversely, if QIH alters the subsequent trajectory itself, functional recovery would demonstrate that neural networks can retain function without preserving their preceding course of evolution. We tested these predictions using epileptogenesis. A return to the preceding network trajectory would leave disease progression largely unchanged, whereas a shift toward a different trajectory could alter the development of electrophysiological and histological pathology. By examining QIH-induced changes in neuronal population activity and their relationship to subsequent network evolution, we sought to identify network properties associated with the divergent functional outcomes of population reorganization.

## Results

### QIH induces a persistent reorganization of hippocampal population activity

We first asked whether hippocampal activity returns to its pre-hibernation state after recovery from QIH. Consistent with our previous observations, tetrode recordings showed that hippocampal spike activity was profoundly suppressed during QIH and recovered after arousal (Fig. 1a,b). Analysis of local field potentials further showed no significant differences in hippocampal oscillatory power before and after QIH across the examined frequency bands (Fig. 1c). Thus, measurements of both neuronal spiking and LFP oscillations indicated an apparent recovery of hippocampal activity to its pre-QIH condition.

**Fig 1.**
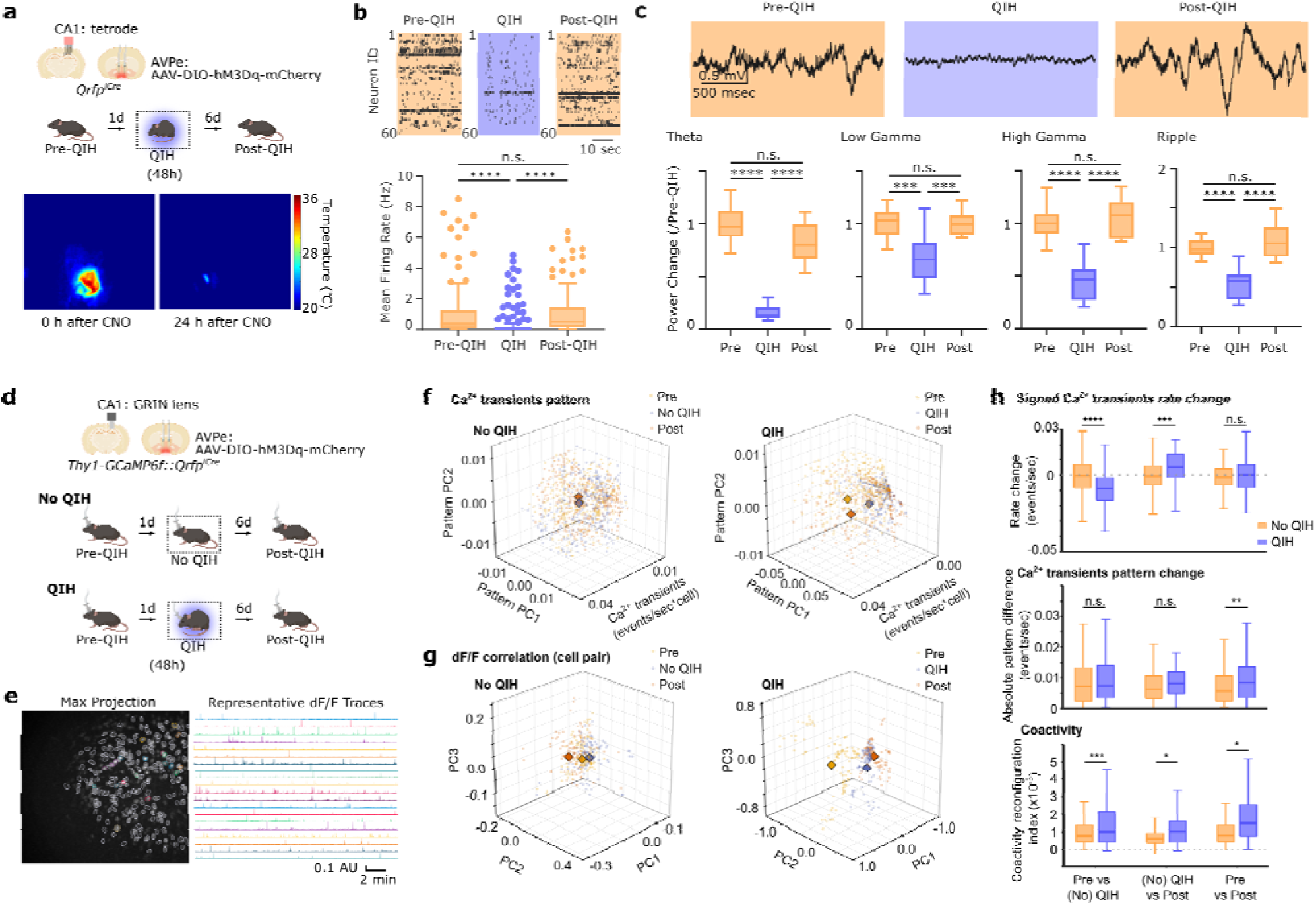
Artificial hibernation induces a hippocampal network state distinct from pre-hibernation despite recovery of overall activity. **a,** Experimental design for electrophysiological recordings during artificial hibernation. QIH was induced by chemogenetic activation of Qrfp-expressing neurons in the anteroventral periventricular nucleus (AVPe), and hippocampal CA1 activity was recorded before QIH, during QIH and after recovery. Representative thermographic images show the reduction in body temperature following QIH induction. **b,** Representative CA1 spike rasters obtained before, during and after QIH (top), and mean firing rates across the three sessions (bottom). Each point represents one recorded unit. Hippocampal firing was strongly suppressed during QIH and recovered after QIH (non-matched units; n = 208 units for Pre-QIH, 141 units for QIH, and 97 units for Post-QIH, Kruskal-Wallis test, p < 0.0001, multiple comparison with Dunn’s correction, QIH vs Pre-QIH, p < 0.0001, QIH vs Post-QIH, p < 0.0001, Pre-QIH vs Post-QIH, p = 0.0626). **c,** Representative CA1 LFP traces and relative power in the theta, low-gamma, high-gamma and ripple-frequency bands before, during and after QIH. Band power was normalized to the corresponding pre-QIH value. QIH broadly suppressed hippocampal oscillatory activity, which recovered after QIH (n = 14∼22 recording channels, Kruskal-Wallis test, multiple comparison with Dunn’s correction, theta: p < 0.0001, QIH vs Pre-QIH, p < 0.0001, QIH vs Post-QIH, p < 0.0001, Pre-QIH vs Post-QIH, p = 0.5327, low gamma: p < 0.0001, QIH vs Pre-QIH, p = 0.0001, QIH vs Post-QIH, p = 0.0002, Pre-QIH vs Post-QIH, p > 0.9999, high gamma: p < 0.0001, QIH vs Pre-QIH, p < 0.0001, QIH vs Post-QIH, p < 0.0001, Pre-QIH vs Post-QIH, p > 0.9999, ripple: p < 0.0001, QIH vs Pre-QIH, p < 0.0001, QIH vs Post-QIH, p < 0.0001, Pre-QIH vs Post-QIH, p > 0.9999). **d,** Experimental design for longitudinal calcium imaging of CA1 neuronal populations in QIH and time-matched No QIH mice. Calcium activity was recorded before the intervention, during QIH or the corresponding middle session, and after recovery. **e,** Representative maximum-projection image showing identified CA1 neurons and representative dF/F traces from the same field of view. **f,** Representative gain–pattern manifolds from No QIH and QIH experiments. Calcium transient counts were calculated in temporal bins and decomposed into a population gain component, representing the mean transient rate, and pattern components describing the relative activity of individual neurons. Each point represents one temporal bin, and diamonds indicate session centroids. For visualization, the 90% of points closest to each session centroid are shown. **g,** Representative coactivity manifolds constructed from vectors of pairwise dF/F correlations. Points and diamonds indicate individual analysis windows and session centroids, respectively. The 90% of points closest to each centroid are shown. **h,** Comparisons of changes in Ca^2+^ transient rate (top), absolute changes in the Ca^2+^ pattern (middle) and coactivity reconfiguration (bottom). Comparisons represent pre-versus middle, middle versus post- and pre-versus post-QIH (or No QIH) sessions (n = 320 cells on average per comparison from three mice; linear-mixed effects models with experimental condition as a fixed effect and mouse as a random intercept. p-values were adjusted using the Holm method across the three session pairs. For signed rate change, adjusted p = 6.22 x 10^−10^, 1.71 x 10^−4^, and 0.316 for the pre-to-middle, middle-to-post, and pre-to-post session pairs, respectively. For Ca^2+^ pattern change, adjusted p = 0.654, 0.691, and 0.0063. For coactivity, adjusted p = 5.41 x 10^−4^, 0.0322 and 0.0322). The middle session corresponds to QIH in QIH experiment and the time-matched recording in No QIH experiment. Cell identity was matched separately for each session pair. Coactivity reconfiguration was defined as the mean cross-validated distance across correlation edges incident on each cell (see Methods). Boxes show the median and interquartile range, whiskers extend to 1.5 times the interquartile range, and points beyond the whiskers indicate outliers. *P<0.05, **P<0.01, ***P<0.001, ****P<0.0001; n.s., not significant.

To track hippocampal population trajectories before, during and after QIH, we performed longitudinal calcium imaging of the same neurons across the three sessions (Fig. 1d). As a control for changes associated with repeated imaging and the passage of time, the mice underwent additional three recording sessions at matched intervals without QIH. We first decomposed population activity into a gain component, reflecting the mean calcium-transient rate across the population, and pattern components, reflecting the relative distribution of activity across matched neurons. Relative to time-matched recordings from the No QIH experiment, QIH produced greater changes in the gain component during the pre-to-QIH and QIH-to-post intervals, consistent with the marked suppression and subsequent recovery of neuronal activity during hibernation (Fig. 1f,h). The pre-to-post gain change did not differ significantly between groups, indicating that the overall population activity rate recovered after QIH. By contrast, the pre-to-post change in the pattern components was significantly greater in the QIH experiment than in the No QIH experiment, whereas no difference was detected during the pre-to-QIH or QIH-to-post intervals. Thus, despite recovery of overall activity, QIH produced a persistent redistribution of activity across neurons that exceeded the change associated with the passage of time alone.

We next asked whether QIH altered the interaction structure of the population. For each time window, we represented population coactivity by the pairwise correlations among simultaneously recorded neurons and projected these correlation patterns into a common low-dimensional space. Coactivity states associated with the pre-QIH, QIH and post-QIH sessions occupied separable regions of this space in each animal (Fig. 1g). Cross-validated distances between session centroids exceeded those obtained after permutation. No QIH also showed time-dependent changes in coactivity, indicating that correlation structure undergoes spontaneous longitudinal drift even in the absence of QIH. The magnitude of post-QIH coactivity reconfiguration nevertheless exceeded that observed over the corresponding interval in No QIH. QIH therefore did not simply reproduce the representational drift associated with repeated recording. Instead, the hippocampal network recovered to a state with normal overall activity but a persistently reorganized coactivity structure. These findings show that recovery from QIH preserves overt hippocampal function without restoring the population configuration present before hibernation.

### QIH reorganizes neuronal interactions and sharp-wave ripple coactivity

To address how the persistent reorganization of hippocampal Ca^2+^ coactivity observed after QIH is achieved, we next examined how spike interactions change during hibernation. We first recorded CA1 spiking activity continuously during the early phase of QIH in naive mice and quantified pairwise interactions between session-matched units using mutual information (Fig. 2a) ^18–20^. Mean firing rates declined markedly after QIH induction, whereas mutual information averaged across all recorded unit pairs remained unchanged (Fig. 2b). The preservation of the population-averaged mutual information despite pronounced suppression of spiking suggested that QIH did not uniformly weaken neuronal interactions.

**Fig 2.**
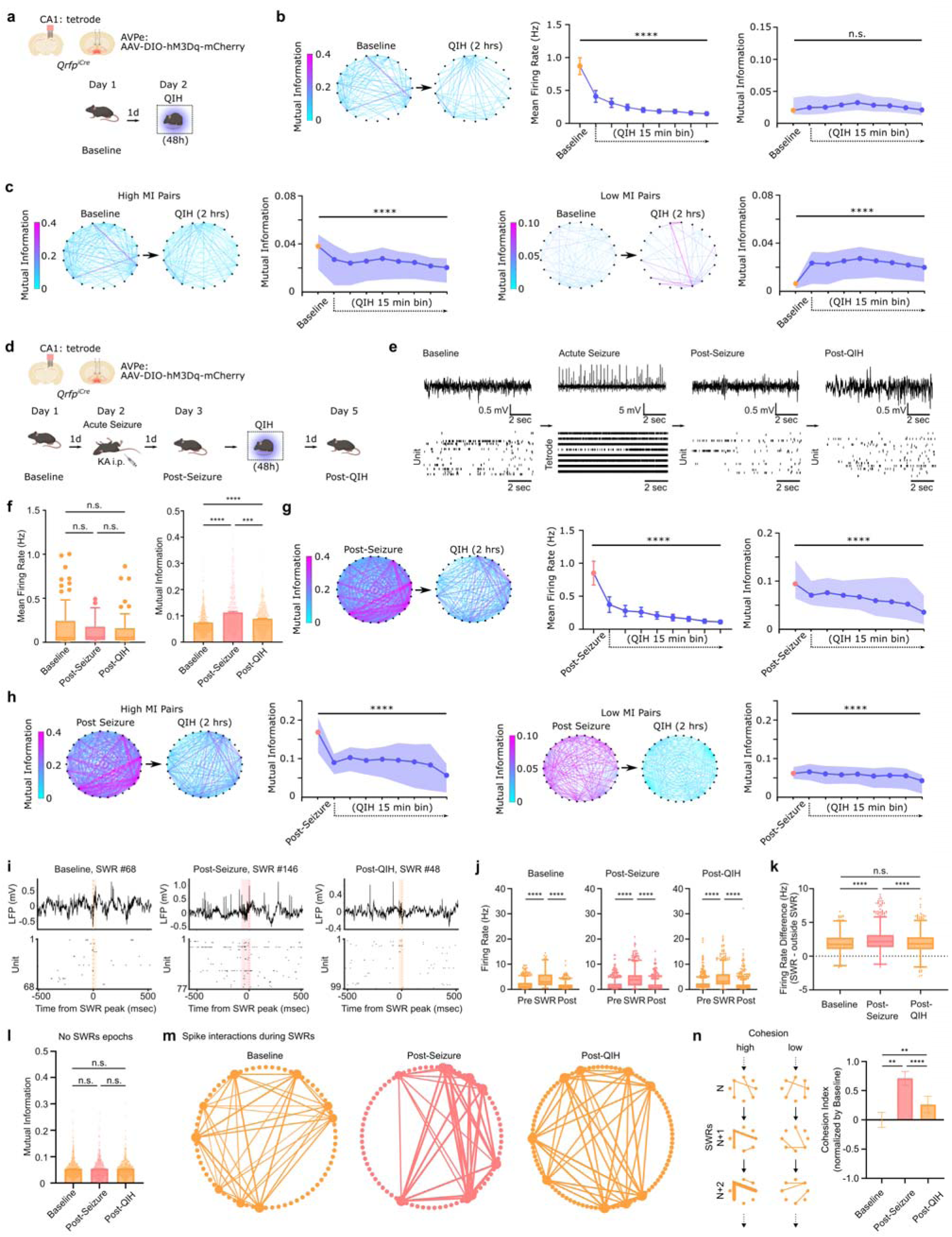
Artificial hibernation reorganizes neuronal interactions in the hippocampus and establishes a distinct sharp-wave ripple coactivity state. **a,** Experimental design for monitoring hippocampal interactions during QIH in naive mice. CA1 unit activity was recorded at baseline and during the initial 2 h of QIH. QIH was induced by chemogenetic activation of Qrfp-expressing neurons in the AVPe. **b,** Representative pairwise mutual information (MI) networks at baseline and during QIH (left). Nodes represent simultaneously recorded CA1 units, and edge color represents MI. Mean firing rate markedly decreased during QIH, whereas population-averaged pairwise MI remained unchanged (right) (n = 121 matched units from three mice; mean firing rate, Kruskal-Wallis test, p < 0.0001; MI, Kruskal-Wallis test, followed by post-hoc multiple comparisons with Dunn’s correction, baseline vs the last 15 min bin during QIH: p = 0.7072). QIH data were divided into consecutive 15-min bins. Lines and shaded regions indicate the median and interquartile range. **c,** MI changes after separating unit pairs according to their baseline MI. Pairs with initially high MI showed a progressive reduction during QIH, whereas pairs with initially low MI showed increased MI (Kruskal-Wallis test, followed by post-hoc multiple comparisons with Dunn’s correction, baseline vs the last 15 min bin during QIH; high MI pairs, left, p < 0.0001; low MI pairs, right, p < 0.0001). Network diagrams show representative changes during the first 2 h of QIH. These opposing changes indicate redistribution of pairwise interactions despite stable population-averaged MI. **d,** Experimental design for examining QIH-induced interaction changes following kainate-induced status epilepticus. CA1 activity was recorded at baseline, after KA-induced seizures, during the initial phase of QIH and after recovery. **e,** Representative CA1 LFP traces and spike rasters at baseline, during an acute seizure, after seizure termination and after QIH. Unsorted population spikes are shown during the acute seizure because individual units could not be reliably isolated during synchronous population discharges. **f,** Mean firing rate of individual units and pairwise MI at baseline, after seizures and after QIH. KA-induced seizures increased pairwise MI without a corresponding persistent increase in mean firing rate, and MI decreased after QIH (Kruskal-Wallis test, followed by post-hoc multiple comparisons with Dunn’s correction; non-matched units, n = 182 units from three mice; firing rate, p = 0.7177 with p > 0.9999 for all three comparisons; MI, p < 0.0001 with p < 0.0001 for Baseline vs Post-Seizure and Baseline vs Post-QIH and p = 0.0001 for Post-Seizure vs Post-QIH). **g,** Representative MI networks immediately before QIH and during the first 2 h of QIH in post-seizure mice. Both mean firing rate and population-averaged MI progressively decreased during QIH (Kruskal-Wallis test, followed by post-hoc multiple comparisons with Dunn’s correction, baseline vs the last 15 min bin during QIH; firing rate, p < 0.0001; MI, p < 0.0001). Values during QIH were calculated in consecutive 15-min bins. **h,** Time courses of initially high- and low-MI pairs in post-seizure mice. Unlike the bidirectional redistribution observed in naive mice, MI decreased during QIH in both pair classes, indicating that the direction of QIH-induced reorganization depends on the preceding network state (Kruskal-Wallis test, followed by post-hoc multiple comparisons with Dunn’s correction, baseline vs the last 15 min bin during QIH; high MI pairs, left, p < 0.0001; low MI pairs, right, p < 0.0001). **i,** Representative sharp-wave ripples (SWRs) and corresponding spike rasters from a mouse at baseline, after seizures and after QIH. **j,** Population firing rates during time windows immediately preceding, overlapping and following individual SWRs at baseline, after seizures and after QIH (Friedman test, followed by post-hoc multiple comparisons with Dunn’s correction; n = 804, 1279, 810 SWRs from three mice; p < 0.0001 for all comparisons). **k,** Within-event increase in population firing rate during SWRs relative to the mean of the matched pre- and post-SWR windows. The SWR-associated firing-rate increase was enhanced after seizures and reduced to baseline levels after QIH (Kruskal-Wallis test, followed by post-hoc multiple comparisons with Dunn’s correction; Baseline vs Post-Seizure, p < 0.0001; Baseline vs Post-QIH, p = 0.5076; Post-Seizure vs Post-QIH, p < 0.0001). **l,** Pairwise MI calculated after removal of all SWR epochs. The post-seizure increase in MI was no longer detected after SWR removal, indicating that SWR-associated activity contributed substantially to the increase in whole-session MI (Kruskal-Wallis test, followed by post-hoc multiple comparisons with Dunn’s correction; Baseline vs Post-Seizure, p > 0.9999; Baseline vs Post-QIH, p = 0.3521; Post-Seizure vs Post-QIH, p = 0.1255). **m,** Representative SWR co-participation graphs at baseline, after seizures and after QIH. Nodes represent all recorded units. Edge weights represent the number of SWRs in which the connected units both fired. **n,** Cross-validated quantification of pair cohesion during individual SWRs (see Methods). Cohesion increased after seizures and decreased after QIH, while remaining distinct from baseline (Linear mixed-effects model with mouse as a random intercept, followed by Holm correction across three pairwise comparisons; Baseline versus Post-Seizure, p = 0.0032; Post-Seizure vs Post-QIH, p = 3.0 x 10^−10^; Baseline vs Post-QIH, p = 0.0032). Boxes show the median and interquartile range, and whiskers extend to 1.5 times the interquartile range. *P<0.05, **P<0.01, ***P<0.001, ****P<0.0001; n.s., not significant.

We therefore examined unit pairs according to their baseline mutual information. Pairs with relatively high mutual information at baseline showed a progressive reduction during QIH, whereas pairs with initially low mutual information showed an increase (Fig. 2c). Thus, the absence of a net change across all pairs reflected opposing changes among different subsets of neuronal interactions. QIH suppressed strong interactions while strengthening initially weak interactions, consistent with a redistribution rather than a uniform loss of hippocampal coactivity.

We next asked how QIH affects neuronal interactions that have already been altered by a pathological perturbation. Mice underwent kainic acid-induced status epilepticus and were recorded at baseline, after the seizure and after arousal from QIH (Fig. 2d,e). Mean firing rates measured across these sessions did not differ significantly. By contrast, pairwise mutual information increased after status epilepticus and declined following QIH to a level that did not differ significantly from baseline (Fig. 2f). To determine whether this decrease reflected the spontaneous evolution of neuronal interactions after status epilepticus, we examined a time-matched cohort that did not undergo QIH (Extended Data Fig. 1a). Mean firing rates remained unchanged, whereas pairwise mutual information increased after status epilepticus and increased further over the subsequent 48-h period (Extended Data Fig. 1b). Elevated neuronal interactions therefore did not spontaneously return toward baseline over the interval corresponding to QIH, but instead continued to increase. Continuous recordings during QIH showed a marked decrease in firing rate accompanied by a progressive reduction in mean mutual information (Fig. 2g). Unlike the opposing changes observed in naive mice, both high- and low-mutual-information pairs decreased during QIH after status epilepticus (Fig. 2h). The direction of interaction reorganization during QIH therefore depended on the preceding network condition. QIH redistributed interactions in the naive hippocampus but broadly reduced the elevated interactions present after status epilepticus.

Because hippocampal sharp-wave ripples are prominent epochs of coordinated population activity, we next examined whether they contributed to the seizure-associated increase in mutual information. We found distinct patterns of unit recruitment during sharp-wave ripples at baseline, after status epilepticus and after QIH (Fig. 2i). Firing rates increased during sharp-wave ripples relative to the immediately preceding and following periods in all three sessions (Fig. 2j). However, the increase in firing rate associated with individual sharp-wave ripples was greater after status epilepticus than at baseline and was reduced after QIH to a level that did not differ significantly from baseline (Fig. 2k). Moreover, when sharp-wave ripple epochs were excluded from the recordings, mutual information no longer differed significantly among the three sessions (Fig. 2l). These results indicate that sharp-wave ripple-associated activity made a major contribution to the increase in neuronal interactions observed after status epilepticus.

Finally, we examined the organization of unit co-participation across sharp-wave ripples. We constructed graphs in which nodes represented recorded units and edge weights reflected the frequency with which pairs of units participated in the same sharp-wave ripple (Fig. 2m). Cross-validated pair cohesion, which quantified whether unit pairs co-participated more consistently than expected from their individual participation probabilities, increased after status epilepticus and decreased after QIH (Fig. 2n). Together, these findings show that QIH reorganizes hippocampal neuronal interactions in a state-dependent manner, interrupts their progressive enhancement after status epilepticus and attenuates the seizure-associated enhancement of coordinated activity expressed during sharp-wave ripples.

### QIH constrains the long-term divergence of neural-state trajectories after status epilepticus

The reorganization of neuronal interactions during QIH raised the possibility that QIH alters the subsequent evolution of hippocampal network activity after status epilepticus. To examine this possibility, we recorded CA1 local field potentials longitudinally from before kainic acid-induced status epilepticus through the following 50-day latent phase (Fig. 3a,b). One group underwent 48 h of QIH early in the latent phase, whereas the other group was recorded over the corresponding period without QIH. Longitudinal recordings indicated that abnormal LFP activity progressively emerged after status epilepticus in mice that did not undergo QIH, whereas hippocampal activity remained comparatively stable throughout the observation period in QIH-treated mice. In a representative untreated mouse, recurrent seizure activity was also captured by simultaneous behavioral video and EEG recordings (Extended Data Video 1).

**Fig 3.**
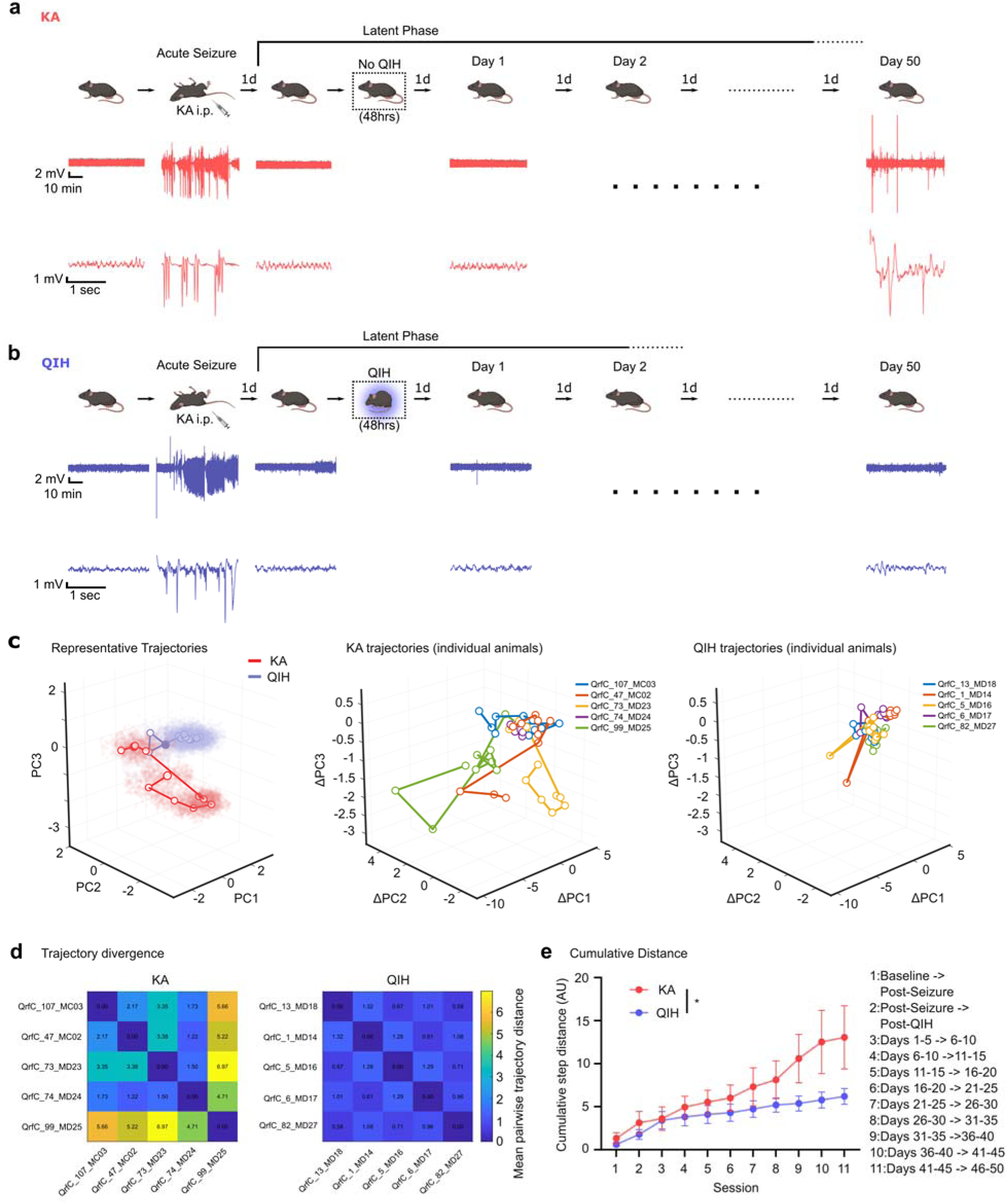
QIH suppresses the long-term divergence of hippocampal network state trajectories after status epilepticus. **a, b,** Experimental design and representative hippocampal LFP recordings from mice subjected t kainic acid-induced status epilepticus followed by either no QIH (a, KA group) or 48 h of QIH (b, QIH group). Hippocampal LFP activity was longitudinally recorded before kainic acid administration, after status epilepticus and throughout the subsequent 50-day latent period. Representative LFP traces are shown at the indicated recording points. Expanded traces are shown below. Scale bars, 2 mV and 10 min for the upper traces and 1 mV and 1 s for the expanded traces. **c,** Principal-component representations of longitudinal hippocampal LFP dynamics. Left, representative neural-state distributions and centroid trajectories from KA and QIH mice. Each point represents the centroid of the LFP-state distribution in one recording session, and successive sessions are connected chronologically. Middle and right, longitudinal trajectories of individual mice in the KA and QIH groups, respectively. Trajectories are expressed as displacement in principal-component space from the first session. KA mice progressively followed animal-specific divergent trajectories, whereas trajectories from QIH mice remained confined to a more restricted region of state space. **d,** Pairwise distances between the longitudinal LFP trajectories of individual animals in the KA and QIH groups. Heat-map values indicate the distance between each pair of animal trajectories. Trajectories became markedly heterogeneous among KA mice but remained comparatively similar among QIH mice. **e,** Cumulative distance travelled through LFP state space across successive recording intervals. Distances were calculated between consecutive recording periods and cumulatively summed for each animal. The numbered transitions correspond to baseline to post-seizure, post-seizure to post-QIH or the corresponding time point in KA controls, and successive 5-day intervals through days 46–50, as indicated (Mixed-effect analysis; N = 5 mice for each group, significant main effect of group, p = 0.0246).

To characterize these longitudinal changes at the level of neural state, we projected LFP activity from all recording sessions into a common principal-component space and tracked the resulting trajectory for each animal (Fig. 3c). In mice that did not undergo QIH, trajectories progressively departed from their initial states and extended in different directions across animals. By contrast, trajectories from QIH-treated mice remained concentrated within a more restricted region of the state space. Thus, status epilepticus was followed by heterogeneous and progressive changes in hippocampal activity, whereas QIH constrained the subsequent range of network-state evolution. We next quantified the divergence of these longitudinal trajectories across animals. Pairwise distances between trajectories were substantially greater in the KA group than in the QIH group, indicating that QIH reduced the inter-animal heterogeneity of network-state evolution (Fig. 3d). We also calculated the cumulative distance travelled through state space across successive recording sessions. Cumulative trajectory distance increased throughout the latent phase in both groups but was significantly smaller in QIH-treated mice (Fig. 3e). Together, these results show that QIH early in the latent phase produces a lasting constraint on the progressive divergence of hippocampal network activity following status epilepticus.

### Divergent neural-state trajectories are accompanied by progressive pathological activity

We next asked whether the divergence of hippocampal state trajectories was accompanied by established electrophysiological features of epileptogenesis. We first quantified pathological high-frequency oscillations (pHFOs) from the longitudinal LFP recordings (Extended Data Fig. 2a) ^21–27^. In mice that did not undergo QIH, the occurrence rate of pHFOs increased during the later stages of the observation period (Fig. 4a). Their power and duration also became progressively elevated relative to QIH-treated mice. By contrast, pHFOs remained comparatively infrequent, low in power and short in duration following QIH. The suppression of trajectory divergence was therefore accompanied by a sustained reduction in pathological high-frequency activity.

**Fig 4.**
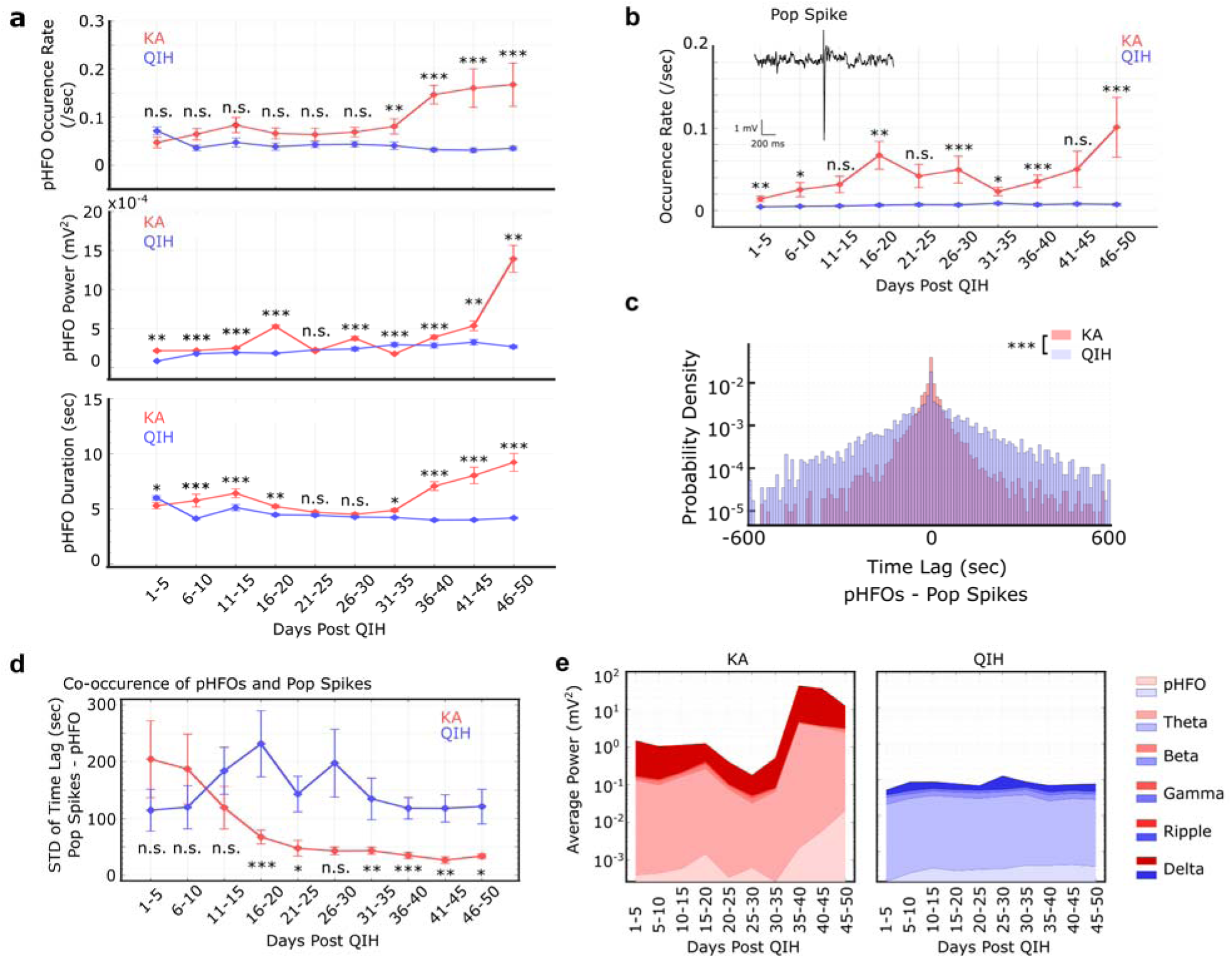
Divergence of hippocampal LFP-state trajectories is accompanied by progressive pathological activity. **a,** Longitudinal development of pathological high-frequency oscillations (pHFOs) following kainic acid-induced status epilepticus. pHFO occurrence rate, mean pHFO power and mean pHFO duration are shown across consecutive 5-day intervals in KA and QIH mice. KA mice showed progressive increases in pHFO occurrence, power and duration, particularly during the later recording periods. These changes were largely absent in QIH mice. **b,** Longitudinal changes in the occurrence rate of population spikes. Inset, representative population spike detected in the hippocampal LFP. Population spikes progressively increased in KA mice but remained infrequent throughout the recording period in QIH mice. Scale bars, 1 mV and 200 ms. **c,** Probability distributions of the temporal lag between pHFOs and population spikes across the longitudinal recordings. The distribution in KA mice was concentrated around zero lag, indicating that the two pathological events increasingly occurred in close temporal proximity. In QIH mice, temporal relationships between pHFOs and population spikes remained less distributed than KA mice. **d,** Temporal dispersion of pHFO–population spike co-occurrence, quantified as the standard deviation of their time lags in each recording interval. The temporal dispersion progressively decreased in KA mice, consistent with the emergence of tightly coupled pathological events, but remained high in QIH mice. **e,** Longitudinal changes in LFP spectral power in the KA and QIH groups. Average band-limited power is shown for delta, theta, beta, gamma, ripple and pHFO frequency ranges across consecutive 5-day intervals. KA mice developed pronounced abnormalities in LFP power during the later stages of epileptogenesis, whereas the spectral profile of QIH mice remained comparatively stable. Group comparisons were performed using the nonparametric rank-sum test, followed by multiple comparisons with Bonferroni p-value corrections. Line plots connect mean values; error bars indicate the standard error of the mean. In each panel, p-values less than 0.05, 0.01, 0.001, 0.0001 were presented as *, **, ***, ****, respectively.

We next examined population spikes, which provide an additional measure of hypersynchronous hippocampal activity (Extended Data Fig. 2b) ^27–29^. Population spikes occurred more frequently in the KA group than in the QIH group across most of the latent phase, with the difference becoming particularly pronounced at later time points (Fig. 4b). In QIH-treated mice, population-spike occurrence remained low throughout the 50-day recording period.

pHFOs and population spikes also became increasingly coordinated in mice that did not undergo QIH, consistent with earlier reports identifying such coupling as a biomarker of epileptic activity ^23–26^. The distribution of temporal offsets between the two event types was more strongly concentrated around zero in the KA group than in the QIH group (Fig. 4c). Consistent with this observation, the variability of their temporal offsets progressively decreased in the KA group, indicating increasingly consistent temporal coupling between pathological high-frequency oscillations and population spikes (Fig. 4d). This temporal coupling did not develop to the same extent following QIH.

Finally, spectral analysis revealed a progressive increase in LFP power in the KA group, with the largest changes emerging during the later stages of the latent phase (Fig. 4e). In contrast, oscillatory power remained comparatively stable across the observation period in QIH-treated mice. Thus, the divergent state trajectories observed after status epilepticus were accompanied by the coordinated emergence of pHFOs, population spikes and abnormal spectral activity. QIH suppressed both the long-term divergence of hippocampal activity and the electrophysiological abnormalities associated with that divergence.

### QIH attenuates the long-term histopathological consequences of status epilepticus

We next asked whether the divergent activity trajectories and progressive electrophysiological abnormalities observed during the latent phase were accompanied by established histopathological features of temporal lobe epilepsy. Following completion of the 50-day LFP recording period, brains were collected and examined for mossy fiber sprouting, neuronal loss and loss of parvalbumin-expressing interneurons.

We first assessed aberrant mossy fiber sprouting using Timm staining (Fig. 5a) ^30^. Mice subjected to status epilepticus without QIH exhibited increased Timm signals in CA3, consistent with the development of mossy fiber sprouting. This increase was not detected in QIH-treated mice, in which normalized Timm signals remained comparable to that in naive controls. Early QIH therefore suppressed the aberrant reorganization of mossy fiber projections that normally develop after status epilepticus.

**Fig 5.**
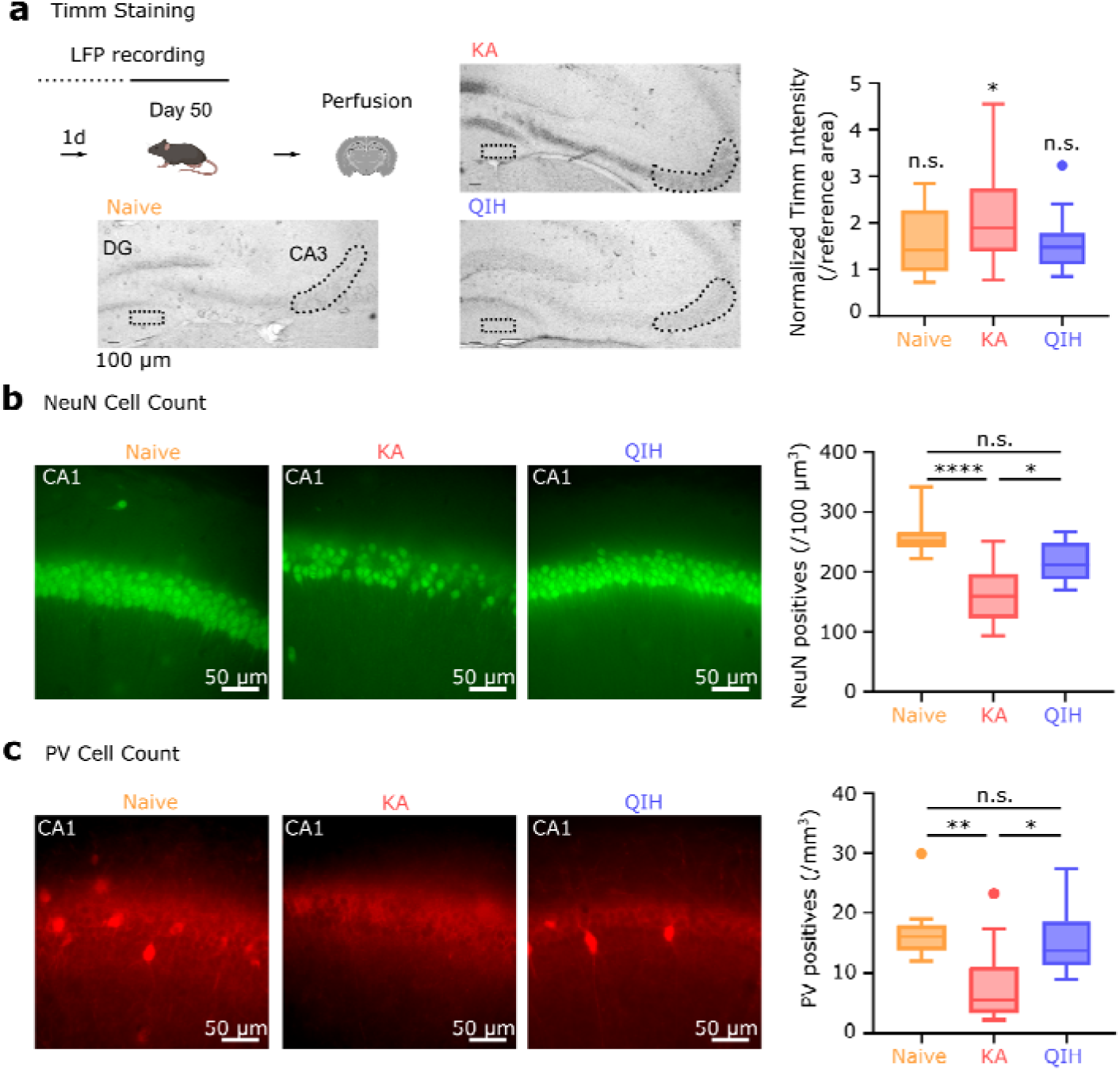
QIH prevents structural hallmarks of epileptic histopathology. **a,** Mossy fiber sprouting assessed after completion of the 50-day longitudinal LFP recordings. Left, experimental timeline and representative Timm-stained hippocampal sections from naive, KA and QIH mice. The outlined regions indicate the area used to quantify aberrant mossy fibre sprouting and the corresponding reference region. Right, Timm staining intensity normalized to the reference area. KA mice exhibited increased Timm staining consistent with mossy fibre sprouting, whereas QIH mice showed no detectable increase relative to naive controls (two-way ANOVA, the main effect of area p = 0.013 and group p = 0.0071, multiple comparisons (CA3 vs the reference area) with Sidak correction, Naïve p = 0.8803, KA p = 0.0406, QIH = 0.1717). Scale bar, 100 μm. **b,** Hippocampal neuronal survival after the longitudinal recording period. Left, representative images of NeuN immunoreactivity in CA1 from naive, KA and QIH mice. Right, density of NeuN-positive cells in CA1. KA mice showed pronounced neuronal loss, which was attenuated in mice that underwent QIH. NeuN-positive cell density in QIH mice was not significantly different from that in naive controls (one-way ANOVA, p < 0.0001, multiple comparisons with Sidak correction, naïve vs KA p < 0.0001, naïve vs QIH p = 0.0664, KA vs QIH p = 0.0182). Scale bars, 50 μm. **c,** Survival of parvalbumin-expressing interneurons. Left, representative images of PV immunoreactivity in CA1 from naive, KA and QIH mice. Right, density of PV-positive cells in CA1. PV-positive interneurons were reduced in KA mice but preserved in QIH mice, with no significant difference between QIH and naive animals (one-way ANOVA, p < 0.0057, multiple comparisons with Sidak correction, naïve vs KA p < 0.0071, naïve vs QIH p = 0.8759, KA vs QIH p = 0.0373). Scale bars, 50 μm.

We next quantified neuronal survival in CA1 using NeuN immunostaining (Fig. 5b). NeuN-positive cell density was markedly reduced in the KA group relative to naive controls. QIH-treated mice retained significantly more NeuN-positive cells than untreated KA mice, and their neuronal density did not differ significantly from that of naive mice. A similar pattern was observed for parvalbumin (PV)-expressing interneurons (Fig. 5c). Their density was significantly higher in QIH mice than KA mice, which did not differ significantly from naive controls.

Thus, the progressive divergence of hippocampal activity and emergence of epileptiform events in untreated mice culminated in mossy fiber sprouting and the loss of both CA1 neurons and PV-expressing interneurons. QIH administered early in the latent phase attenuated each of these long-term histopathological abnormalities. Together with the longitudinal electrophysiological results, these findings indicate that QIH redirects the post-seizure network away from a trajectory associated with progressive functional and structural pathology.

## Discussion

Our findings show that recovery from a profound physiological perturbation does not require restoration of the preceding neural state. Although hippocampal activity recovered after QIH, the population entered a distinct state characterized by persistent reorganization of neuronal activity patterns and coactivity. The form of this reorganization depended on the state of the network before QIH. In the naive hippocampus, QIH redistributed interactions among neurons without producing a net loss of population-wide interaction strength. By contrast, when the network had been altered by status epilepticus, QIH reduced the elevated neuronal interactions and excessive coactivity expressed during sharp-wave ripples. This early reorganization was followed by constrained divergence of hippocampal LFP trajectories and attenuation of the electrophysiological and structural abnormalities that otherwise developed over the subsequent 50 days. Together, these findings support a model in which function-preserving and pathological state changes are distinguished not by the magnitude of neural reconfiguration or by proximity to a preceding state, but by the organization of neuronal interactions and the subsequent network trajectories associated with that organization (Fig. 6).

**Fig 6.**
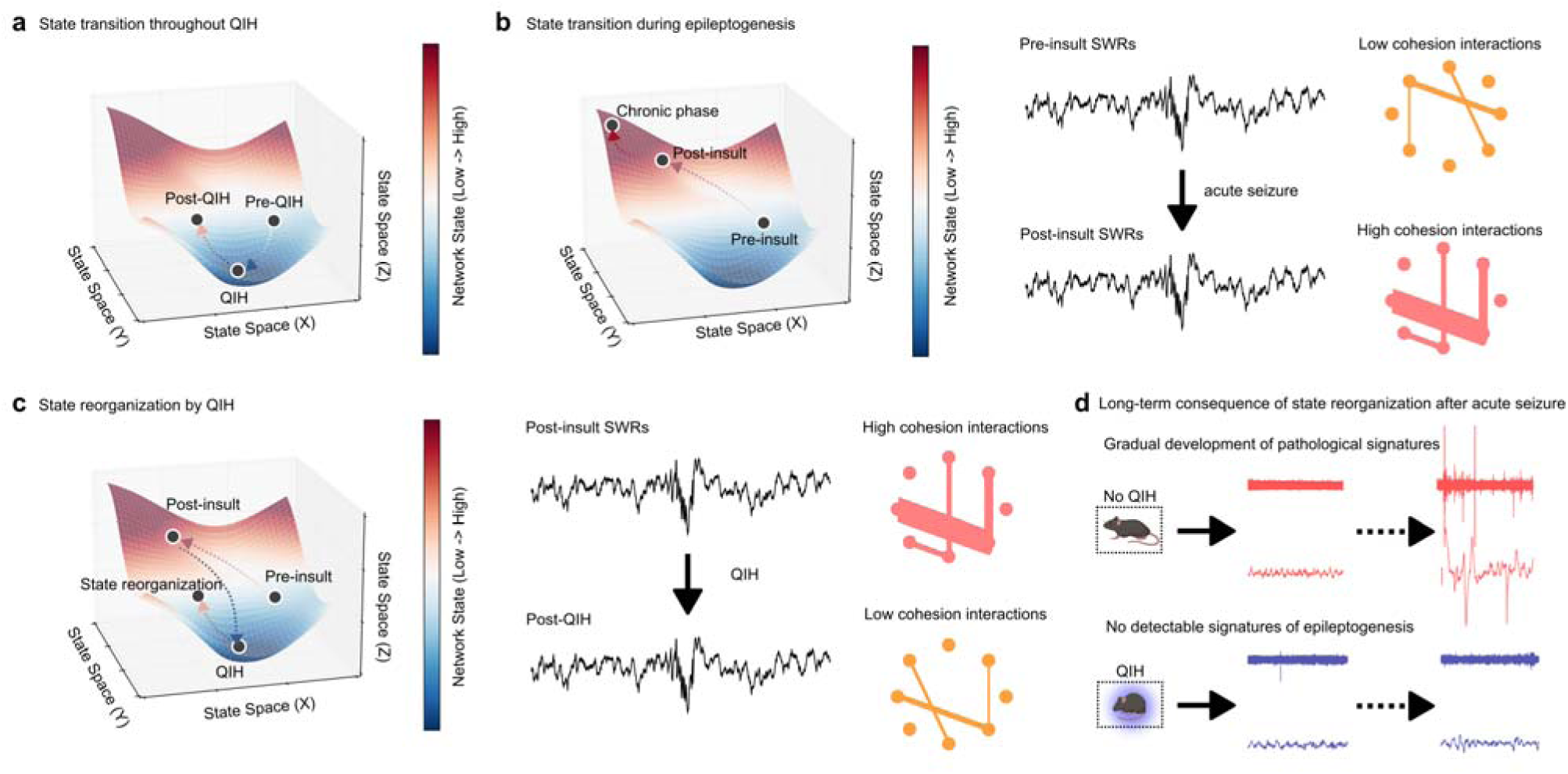
Working model for QIH-induced reconfiguration of hippocampal network states and suppression of epileptogenesis. **a,** QIH induces a transition between distinct hippocampal network states. Neural activity moves from the pre-QIH state into an extreme QIH-associated state and subsequently enters a post-QIH state that is distinct from both the pre-QIH and QIH states. Thus, arousal from QIH does not simply restore the preceding network configuration. **b,** Network-state progression during epileptogenesis. Acute seizures alter hippocampal neuronal interactions, including increased coactivity and cohesion during sharp-wave ripples. The post-insult network progressively diverges from its initial state and develops chronic electrophysiological and histological features of epilepsy. **c,** Reconfiguration of the post-insult network by QIH. QIH transiently moves the hippocampal network into a markedly different state and reorganizes neuronal interactions during sharp-wave ripples. After arousal from QIH, the network enters a new state characterized by reduced SWR-associated coactivity rather than returning directly to the post- or pre-insult state. This transition redirects the network away from its otherwise pathological trajectory. **d,** Long-term consequences of QIH-induced state reconfiguration. Without QIH, pathological electrophysiological signatures progressively emerge and are followed by chronic hippocampal pathology. When QIH is induced during the latent phase, the pathological network-state trajectory is constrained, and detectable electrophysiological and histological signatures of epileptogenesis are largely absent over the subsequent observation period.

The finding that hippocampal activity does not return to its pre-QIH population configuration refines, rather than contradicts, our previous conclusion that memory-related representations survive artificial hibernation ^17^. Our previous work showed that memories acquired before QIH remained behaviorally accessible after arousal and that the quality of hippocampal spatial coding was preserved. Tetrode recordings revealed no detectable deterioration in place-field properties or population decoding accuracy after QIH, while longitudinal calcium imaging showed that changes in place fields across QIH were no greater than the representational drift observed over the same period without QIH. These measurements established that QIH does not disrupt the spatial information carried by hippocampal populations, but they did not determine whether the same information was supported by an identical population configuration. By examining the distribution of activity across matched neurons and the relationships among simultaneously recorded cells, the present study revealed changes that were not apparent from place fields or decoding performance alone. Overall activity recovered, yet its allocation across neurons and the underlying coactivity structure were reorganized after QIH. The same representational content can therefore be maintained without reconstructing the precise neuronal relationships that existed before hibernation. This interpretation is also consistent with the structural observations of our previous study. Although many dendritic spines lost during QIH reappeared near their original locations after arousal, this apparent spine recovery did not account for memory preservation, and spine imaging could not establish whether the recovered structures reconnected with the same presynaptic partners. The altered coactivity structure observed here does not by itself demonstrate synaptic rewiring, but it suggests that recovery of spine number and morphology need not entail restoration of the original connection pattern. QIH-induced reconfiguration should therefore not be understood as wholesale replacement of a hippocampal representation. Rather, it reflects a selective reorganization of neuronal relationships within a network that retains its established representational content and memory function.

The state dependence of this reorganization distinguishes QIH from a simple global scaling process. Homeostatic synaptic scaling is classically understood as a compensatory mechanism in which prolonged changes in neuronal activity induce coordinated increases or decreases in synaptic strength, often through proportional changes that preserve the relative strengths of individual synapses ^12,13,31^. Although pairwise mutual information does not directly measure synaptic efficacy, an analogous interaction-level scaling process would be expected to shift neuronal interactions predominantly in the same direction or to preserve their relative organization. This was not observed during QIH. In the naive hippocampus, the marked reduction in firing was accompanied by weakening of initially strong interactions and strengthening of initially weak interactions, leaving population-averaged mutual information unchanged. Following status epilepticus, the same physiological perturbation instead reduced interactions across both initially high- and low-mutual-information pairs. QIH therefore did not produce a fixed interaction state or a uniform response to reduced activity. Rather, the direction of reorganization depended on the organization of the network before QIH, suggesting that QIH engages a state-dependent capacity to redistribute neuronal interactions.

SWRs provide a potential functional context for linking neuronal interactions to pathological progression. These highly synchronous hippocampal events coordinate cell-assembly reactivation and communication with distributed cortical networks during memory consolidation and retrieval ^32–35^. In hyperexcitable networks, however, the circuitry that normally supports coordinated neuronal recruitment during SWRs can also give rise to pathological synchronization, potentially facilitating the amplification and propagation of epileptiform activity ^36–39^. Epileptogenesis is increasingly understood as a network process in which abnormal activity arising within a restricted region recruits anatomically connected circuits ^11,40,41^. Consistent with this view, hippocampal interictal discharges can drive cortical oscillations with temporal coupling that exceeds physiological ripple– spindle coupling, suggesting that pathological activity can engage communication pathways normally involved in hippocampal–cortical coordination ^42^. Our results do not establish that the altered SWRs observed after status epilepticus propagate pathological activity to other regions. They nevertheless show that individual SWRs recruit more firing and more consistently coactive neuronal pairs after status epilepticus, creating a population event with greater potential to amplify and distribute locally generated activity. QIH reduced both the excess firing and pair cohesion expressed during SWRs without abolishing SWR-associated activity. QIH may therefore reduce the reinforcement and propagation of local hyperactivity by limiting excessive neuronal recruitment during SWRs, while preserving the physiological event itself.

Several questions remain unresolved. Our recordings focused primarily on CA1, and the contributions of other hippocampal subregions and connected cortical structures to QIH-induced reorganization remain to be determined. More importantly, hibernation simultaneously alters body temperature, metabolism, neuromodulatory tone and synaptic structure ^43–45^. The present study therefore cannot identify which of these changes is necessary or sufficient to reorganize neuronal interactions and constrain subsequent network evolution. Separating these components will be important for establishing whether the protective effect requires the complete QIH state or can be reproduced by manipulating a more specific cellular or circuit mechanism. Finally, we induced QIH early after status epilepticus, before the progressive electrophysiological and structural abnormalities examined here had become established. Whether a comparable state transition can modify network trajectories after chronic epilepsy has developed remains unknown. More broadly, it will be important to determine whether QIH can alter maladaptive network reorganization following other forms of brain injury or disease.

Together, our findings suggest that the functional consequences of neural population reorganization cannot be inferred from the extent of changes in population configuration alone. QIH profoundly reorganized neuronal interactions while attenuating the progressive changes in network activity and pathology associated with epileptogenesis. The significance of a neural population state may therefore lie not only in the information it represents at a given moment, but also in how its organization relates to the subsequent evolution of network activity. The course of network evolution can be redirected through profound yet reversible physiological transitions, such as hibernation.

## Methods

### Animals

Qrfp-iCre mice were generated and maintained on a C57BL/6J background, then crossed with CD1 mice to obtain F1 hybrid animals carrying the Qrfp-iCre gene. These single transgenic mice were used for electrophysiology experiments. As naïve animals for Figure 3, C57BL/6J WT mice were used. For Ca^2+^ imaging, Thy1-GCaMP6f (JAX 024276) mice were crossed with Qrfp-iCre mice (C57BL/6J background) to obtain F1 hybrid animals carrying both transgenes. Both males and females were randomly assigned to each of the experimental groups in the study. All animals were provided water and food ad libitum and maintained in a temperature- and humidity-controlled room with a 12 h light/dark cycle (lights on from 07:00 A.M. to 7:00 P.M.) in accordance with AAALAC accreditation standards. The OIST Animal Care and Use Committee approved all experimental protocols.

### Surgery

AAVs were either purchased from Addgene or produced by using AAVpro Purification Kit (Takara, Japan). Titers of the vectors were as follows: AAV.hsyn.DIO.hM3D(Gq).mCherry, 2.30 x 1013; AAV.hSyn.DIO.mCherry, 1.70 x 1013 (copies/mL).

All animals were single-housed after surgery. Mice were anesthetized with i.p. injection of MMB (a mixture of midazolam, medetomidine, and butorphanol tartrate at concentrations of 4.0 mg/kg, 0.3 mg/kg, and 5.0 mg/kg, respectively). For hM3Dq (or control mCherry) expression in Q neurons of AVPe (AP, 0.38 mm; ML, ±0.3 mm; DV, −5.1 mm), 300 μL of AAV-DIO-hM3Dq-mCherry or AAV-DIO-mCherry was injected. More specifically, each AAV was infused through a small craniotomy above the site at a rate of 100 μL per minute with a 33G blunt needle (NF33BL, NanoFil) and a micropump syringe (World Precision Instrument). After infusion, the 33G needle stayed at the infusion site for 5 minutes to allow diffusion of the vector.

The microdrives for single-unit and LFP (Local Field Potential) recording were constructed as previously described 42. It had eight independently adjustable tetrodes (14 nm diameter, nichrome wire, gold-plated to an impedance of 200-300 kΩ) and another adjustable tetrode for reference. The Microdrive was implanted to target the dorsal CA1 (AP, −2.0 mm; ML, 1.5 mm; DV, −1.3 mm) for single unit recording or the intermediate part of CA1 (AP, −3.0 mm; ML, 3.0 mm; DV, −2.2 mm) for LFP recording. One of the stainless screws attached to the skull was used as an electrical ground for extracellular recording. After recovery, the depth of the tetrodes was slowly and manually adjusted until reliable detections of sharp wave ripples were achieved for LFP recording. For spike recording, tetrodes were further adjusted until multiple spike clusters appeared. During the adjustments, mice stayed in a highly familiar small plastic bucket (20 cm diameter) located in the recording room.

For one-photon Ca2+ imaging, GRIN lens (ProView Integrated Lens, 1.0 mm diameter, 4.0 mm lens length, Inscopix, USA) was implanted at the dorsal CA1 of the hippocampus (AP, −2.0 mm; ML, 1.5 mm; DV, 1.6 mm). Cortical tissue above the target corpus callosum was aspirated using 26G and 27G needle to secure the lens path.

For EEG recording, the EEG headmount (Pinnacle Technology, 8201) was glued to the surface of the skull, and screws (Pinnacle Technology, 8209) connecting the headmount were implanted into the craniotomy hole created by a surgical drill.

### QIH induction

Before QIH induction, mice were handled by an experimenter for at least 5 days. Two to three weeks after surgery, 0.1 mL/g of saline was injected intraperitoneally (i.p.) once daily for 4 days to habituate the i.p. injection. One day before the QIH injection, the mice were shaved on the back to measure body temperature at the surface.

On the day of QIH induction, mice were allowed to habituate in the hibernation chamber (HC-10, Shin Factory) for 1 hour. The chamber is air-circulated, set at 20 °C, and under constant darkness. Within the hibernation chamber, each animal was placed in a black plastic box (9 x 18 x 20 cm, no ceiling) with bedding materials, water gel, and food pellets provided. Their body temperature was recorded using the infrared (IR) thermal camera (FLIR C5) attached 50 cm above the box. The temperature and location of the animal were identified and tracked by using custom-made software (HIBER: https://github.com/oist/Tanaka_K_Lab_public).

After habituation, CNO (1 mg/kg) was i.p. injected to activate hM3Dq-expressing Q-neurons in AVPe. Mice were allowed to stay in the hibernation chamber for 48 hours, during which their temperatures and locations were recorded at 1 Hz using the IR camera and HIBER. After 48 hours, the mice were returned to their home cage in the vivarium and allowed to recover without disturbance.

### Kainic acid-induced status epilepticus

Prior to acute seizure induction, mice were placed in a highly familiar box and connected to a 32-channel Digital Lynx SX acquisition system for LFP recording. The baseline LFP was first recorded for 15 minutes to ensure reliable recording and apparent normal activity. Then, for acute seizure induction, 20 mg/kg of kainic acid (KA; ab120100, Abcam, USA) dissolved in saline was i.p. injected while monitoring LFP signals. Successful seizure induction was confirmed through the high-amplitude epileptic burst activity. These hyperactivities indicative of acute seizures were monitored for 2 hours after KA injection, and then the mice were disconnected from the cable for recovery in the vivarium. Mice that did not survive the acute seizure were excluded from the experiments.

### Tetrode recording

All tetrode data were acquired using a 32-channel Digital Lynx SX acquisition system (Neuralynx). LFP was filtered at 2-9000 Hz and down-sampled to 1600 Hz to reduce data size while preserving the frequency range of interest for high-frequency oscillations (0 to 800 Hz). Spike waveforms were filtered between 0.6-6 kHz, and those above a peak threshold of 50 μV were time-stamped and digitized at 32,556 Hz.

For unit recording at a resting state, mice were placed in a familiar box (17 cm in diameter) and recorded for 15 minutes.

After the acute seizure with or without subsequent QIH induction, LFP recording was conducted for 1 hour once a day for 50 days. Each day, mice were first placed in a familiar box and left for at least 30 minutes to minimize the effects of transportation. A 1-hour LFP recording was conducted following confirmation of LFP signatures in the hippocampus.

After all recording sessions, mice were anesthetized for electrolytic lesions to mark tetrode locations and then transcardially perfused. Following 24 hours of post-fixation with 4% PFA (paraformaldehyde), brains were sectioned (50 μm thickness) to confirm tetrode locations in the hippocampal CA1. From each animal, a single tetrode, confirmed to be located in the target based on LFP prior to the experiment and to remain in the target based on LFP on Day 50 and lesion, was used for a 50-day LFP analysis. The brain sections containing the dorsal hippocampus from LFP-recorded mice were collected for immunostaining.

### One-photon Ca^2+^ imaging

We recorded Ca2+ dynamics of the CA1 pyramidal cells in the dorsal hippocampus using an integrated miniature fluorescence microscope (nVista 2.0, Inscopix). Qrfp-iCre::Thy1-GCaMP6f mice were injected with AAV-DIO-hM3Dq-mCherry and received implantation of the GRIN lens targeting the CA1 (AP, −2.0 mm; ML, 1.5 mm; DV, −1.6 mm). Three weeks after surgery, their Ca2+ activities were recorded by using the Inscopix acquisition software with a frame rate of 20 Hz when mice were placed in a familiar box (32.5 cm x 44.5 cm). Ca2+ transients were extracted as delta F/F values higher than 3*standard deviations above the mean. Events with inter-event-intervals of 150 msec or less were combined as a single event.

### EEG recording

Weeks after recovery from the acute seizure induced by KA injection (see above), mice were placed in a clear acrylic glass bucket (Pinnacle Technology, 8228) with bedding materials and ad libitum food gel provided. The mice were connected to a pre-amplifier (Pinnacle Technology, 8202), and the data were acquired by Sirenia Acquisition (Pinnacle Technology, version 2.2) at a sampling rate of 2,000 Hz. Experiment started after habituation to the recording bucket. EEG/EMG signals, and behavioral video acquired from a USB camera (30 fps) were acquired for 24 hours continuously.

### Spike sorting

Spikes were manually sorted using SpikeSort3D software (Neuralynx) as previously described 42. The waveform variables (peak amplitudes and their energies) of all spikes recorded from a combination of three channels were projected to 3D space, and putative units were manually clustered. This procedure was repeated for all combinations of channels. Clusters with > 0.5 % spikes having an inter-spike-interval of < 2 msec, a total number of spikes < 50, or an isolation distance < 10 were excluded. The identified units were not matched across sessions.

### Power analysis

Power spectral density (PSD) was computed using Welch’s method on 4-s windows. Power of specific frequency bands was obtained by integrating PSD values over the following ranges: Delta (2–4 Hz), Theta (4–12 Hz), Beta (12–30 Hz), Gamma (30–100 Hz), sharp-wave ripple (SWR; 100–250 Hz), and pathological high-frequency oscillations (pHFO; 300–800 Hz). Total power was calculated across the full 2–800 Hz range.

### pHFO event detection

HFO events were detected from LFP traces resampled at 1.6 kHz using spectral power analysis. LFP signals were segmented into 4-s windows with 1-s overlap. For each window, band-specific power was calculated by integrating the PSD within predefined frequency ranges as described above. To minimize contamination by line noise, frequencies within ±7.5 Hz of 60 Hz harmonics (up to 800 Hz) were excluded. Windows in which HFO power exceeded the median plus two times the median absolute deviation (MAD) of the HFO power distribution were identified as HFO events.

### Population spikes event event detection

Population (pop) spikes were identified from the LFP signals. The algorithm was tested on a couple of 1-hour example traces, each with population spikes manually tagged by 2 experimenters. Parameters were adjusted so that the procedure identifies most of the tagged population spikes, relying as much as possible on data statistics rather than hardcoded numbers. The selection of the pop spikes events was:

1. Identified periods with sudden larger signal variation. Specifically, when the mean derivative of the signal over a 625 us period was more than five standard deviations (5σ) from the mean.
2. A second pass confirmed these events by ensuring their amplitude was also greater than five standard deviations from the mean.

These computations were performed on sliding 4-second windows to ensure that the standard deviation was representative of the local brain state, thereby minimizing false positives caused by overall increases in activity during particular periods.

### Mutual Information

Spike trains from each cell in each 15-minute recording session were converted into spike count vectors (bin size: 1 second). Mutual information (MI) was computed as the sum of all joint probability distributions as follows:

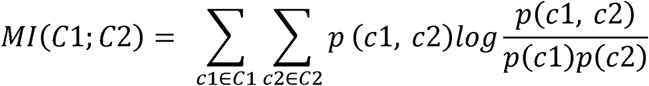

where p(c1, c2) is the joint probability distribution, and p(c1) and p(c2) are the marginal probability distributions of spike count vectors from cell #1 and #2, respectively.

### Gain and activity-pattern decomposition

Calcium transients were binned in time and converted to event rates for each neuron. For each time bin, population activity was decomposed into a gain component, defined as the mean event rate across neurons, and an activity-pattern component, defined as the neuron-specific deviations from this population mean. Pattern vectors were subjected to principal component analysis after concatenating data from the three sessions. Population states were visualized using the gain axis and the first two pattern principal components. For visualization across all sessions, only neurons identified in all three sessions were included. Quantitative session comparisons were performed separately for each pair of sessions using all neurons matched between those sessions.

### Coactivity analyses in Ca^2+^ data

To assess neuronal interactions independently of overall activity, pairwise correlations in calcium activity were calculated within successive time windows. Each window was represented by the vector of correlations across all neuron pairs, and these vectors were concatenated across sessions before principal component analysis. Session separation was quantified using cross-validated distances between coactivity states. Analyses were performed separately for each mouse, and group statistics gave equal weight to each mouse.

### Sharp-wave ripple detection and event-level analyses

Sharp-wave ripples (SWRs) were detected using ripple-band (100-180 Hz) LFP activity together with multi-unit activity recorded from the corresponding tetrode. Candidate events were identified from the envelope of the ripple-band signal and retained when accompanied by increased multi-unit firing. Events adjacent to discontinuities in the LFP time series were excluded. The same detection parameters were applied to all sessions. For each SWR, firing rate was calculated within the detected event and compared with firing rates in duration-matched intervals immediately before and after the event.

### SWR co-participation and graph analysis

For each mouse and session, unit participation was represented as a binary matrix indicating whether each unit fired at least once during each SWR. Co-participation graphs were constructed with units as nodes and edge weights defined by the number of SWRs in which the corresponding unit pair was jointly active.

Building on established approaches that quantify pairwise neuronal coactivity relative to that expected from individual participation probabilities ^46,47^, we computed pair cohesion by using fivefold temporal cross-validation. SWRs were ordered by occurrence time and divided into five contiguous folds. For each fold, the remaining SWRs were used to estimate the participation probability of each unit, *p_i_*, and the joint participation probability of each unit pair,*p_ij._* The affinity of each pair was defined as

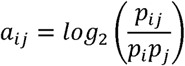

For each held-out SWR containing at least two active units, observed cohesion was calculated as the mean training-derived affinity across all pairs of units participating in that event. A null distribution was generated from 1,000 random sets containing the same number of units, sampled without replacement according to the training-derived participation probabilities.

Cohesion index was calculated as

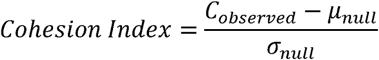

where

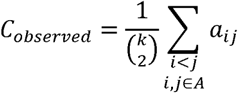

and *μ*_null_ and *σ*_null_ are the mean and standard deviation of the corresponding null distribution. Positive values indicate that the units participating in an SWR had a greater tendency to co-participate during the training events than expected from their individual participation probabilities.

### Longitudinal LFP feature extraction

LFP signals down-sampled at 2,000 Hz were divided into 4-s windows with 1-s overlap, yielding a 3-s step between consecutive windows. Each window was linearly detrended before feature extraction. Power spectral density was estimated using Welch’s method with 1-s Hann segments, 50% overlap and 4,096-point Fourier transforms.

Ten features were used for principal component analysis. These comprised the logarithm of the median absolute deviation of LFP amplitude, the logarithm of line length, Hjorth complexity, log-transformed theta-to-delta power ratio, log-transformed gamma-to-delta power ratio, log-transformed pathological high-frequency oscillation-to-delta power ratio, spectral entropy, spectral flatness, the 95% spectral edge frequency and the slope of the power spectrum in log-frequency–log-power coordinates. Delta, theta, gamma and pathological high-frequency oscillation power were calculated over 2–4 Hz, 4–12 Hz, 30–100 Hz and 300–800 Hz, respectively. Spectral-shape features were calculated over 2–800 Hz after excluding frequencies within ±1 Hz of 60-Hz harmonics.

### LFP population-state manifold and trajectory analyses

Window-level feature vectors from all animals and recording sessions were projected into a common principal-component space. To prevent animals or time points containing larger numbers of windows from dominating the PCA fit, a maximum of 3,000 windows was sampled from each animal and experimental time point. Within this balanced fitting set, each feature was winsorized at the 0.5th and 99.5th percentiles and robustly standardized by subtracting its median and dividing by 1.4826 times its median absolute deviation. PCA was fitted to these standardized features. All valid windows, without winsorization, were then transformed using the same centering, scaling and PCA coefficients.

The first three principal components defined the LFP population-state space. For each mouse and experimental time point, the session state was represented by the median coordinates of all windows along PC1–PC3. Session centroids were ordered chronologically to generate an individual longitudinal trajectory. For visualization, centroid coordinates were expressed relative to the earliest recorded session from the same mouse.

Movement through state space was quantified as the Euclidean distance between successive session centroids. Cumulative trajectory distance was obtained by summing the distances between consecutive centroids across recording periods. Inter-animal trajectory divergence was quantified by comparing the longitudinal centroid trajectories of pairs of mice in the common PC1–PC3 space.

### Histology and immunohistochemistry

The coronal brain sections (50 μm thickness) were first washed with PBS for 5 minutes, three times. The sections were then incubated in blocking buffer (3% goat serum & 0.3% Triton X-100 in PBS) for 15 minutes.

After blocking, the sections were incubated in primary antibody solution at 4 °C overnight (anti-Parvalbumin, ab277625, abcam, 1:1000; anti-NeuN, ABN91, Merck, 1:3000). After the primary incubation, the sections were washed with PBS for 5 minutes three times, followed by incubation in secondary antibody solution for 2 hours at room temperature (anti-chicken, AF555, Invitrogen, A21437; anti-mouse, AF633, Invitrogen, A-21050). After three additional PBS washes for 5 minutes, the sections were mounted on slide glasses using mounting media containing DAPI (ProLongTM Gold Antifade mountant with DAPI, Thermo Fisher).

### Timm staining and mossy fiber sprouting quantification

Timm staining was performed using the FD Rapid TimmStain KitTM (FD NeuroTechnologies, Inc.). Briefly, coronal brain sections (50 μm thickness) containing the dorsal hippocampus were first washed three times with PBS (pH 7.4), then stained with the staining solution for 50 minutes at 30 °C in the dark. The sections were protected from light exposure in the following steps. After staining, sections were washed in Milli-Q for 3 minutes, in running water for 30 minutes, and then in distilled water again for 3 minutes. The sections were then dehydrated in 50%, 75%, 95%, and 100% ethanol (3 minutes each). After dehydration, the sections were cleared in xylene for 3 minutes, and mounted with mounting medium.

Timm-stained hippocampal sections were imaged using a Leica Thunder imaging system under standardized acquisition parameters to ensure consistency across samples. Captured images were processed using Fiji (ImageJ, NIH) and converted to 16-bit TIFF format for subsequent analysis.

For quantitative assessment of Timm staining intensity, regions of interest (ROIs) were selected in the CA3 region. To ensure reproducibility, a standardized ROI template (area = 8007.729 μm²) was applied across all samples using the ROI Manager function in Fiji. The spatial calibration was set to 0.8625 pixels/μm. Mean grayscale intensity values were extracted from each ROI and normalized to those from reference areas (Figure 3A).

### Statistical analysis

Statistical analyses were performed using MATLAB (2024b, 2025a, MathWorks), R, and GraphPad Prism (Version 10, GraphPad Software). Data loading was managed using the Neuralynx API (NlxAPI), and analyses were performed using custom scripts. Data in the main text and figures are represented as mean ± the standard error of the mean, or as boxplots indicating the median, 25th, 75th, the maximum/minimum value within 1.5x the interquartile range, and the outliers, unless otherwise indicated. The significance p-value was set at 0.05 or below. In each figure, p-values less than 0.05, 0.01, 0.001, 0.0001 were presented as *, **, ***, ****, respectively.

## Acknowledgements

We would like to thank Thomas J. McHugh at RIKEN, Joshua P. Johansen at RIKEN, Yoko Yazaki-Sugiyama at OIST, and Inna Slutsky at Tel Aviv University for comments and discussions on an earlier version of the manuscript. We also appreciate all the members of the Memory Research Unit and the supporting sections in OIST. We especially thank K. Nakamura for daily assistance with our research activities, the Animal Resource Section for the maintenance of our mouse colony, and the Engineering Section for 3D printing Microdrive parts. We also thank Arthur J. Y. Huang at RIKEN for DNA plasmids, AAV vectors, and technical advice on AAV production. Some illustrations in figures are created with BioRender.com. This work is supported by OIST institutional research budget (KZT), JSPS KAKENHI Grant-in-Aid for Scientific Research (B) (Grant Number: 21H02585, KZT), Grant-in-Aid for Transformative Research Areas (A) (Grant Number: 23H04944 & 23H04939, KZT), and CREST (Grant Number: JPMJCR24T4, KZT).

## Author contributions

Conceptualization, K.Z.T.; Methodology, D.M., M.C.C., V.M.S., Y.L., H.A., A.T.N., K.Z.T.; Investigation, D.M., M.C.C., V.M.S., Y.L., H.A., A.T.N., K.Z.T.; Formal Analysis, D.M., M.C.C., Y.L., H.A., M.O. K.Z.T.; Visualization, D.M., K.Z.T.; Funding acquisition, K.Z.T.; Project administration, K.Z.T.; Resources, D.M., M.C.C., V.M.S., Y.L., H.A., A.T.N., K.Z.T.; Supervision, K.Z.T.; Writing – original draft, K.Z.T.; Writing – review & editing, D.M., M.C.C., V.M.S., Y.L., H.A., A.T.N., K.Z.T.

## Additional information

Supplementary information is available for this paper.

---

**Extended Data Fig. 1.**
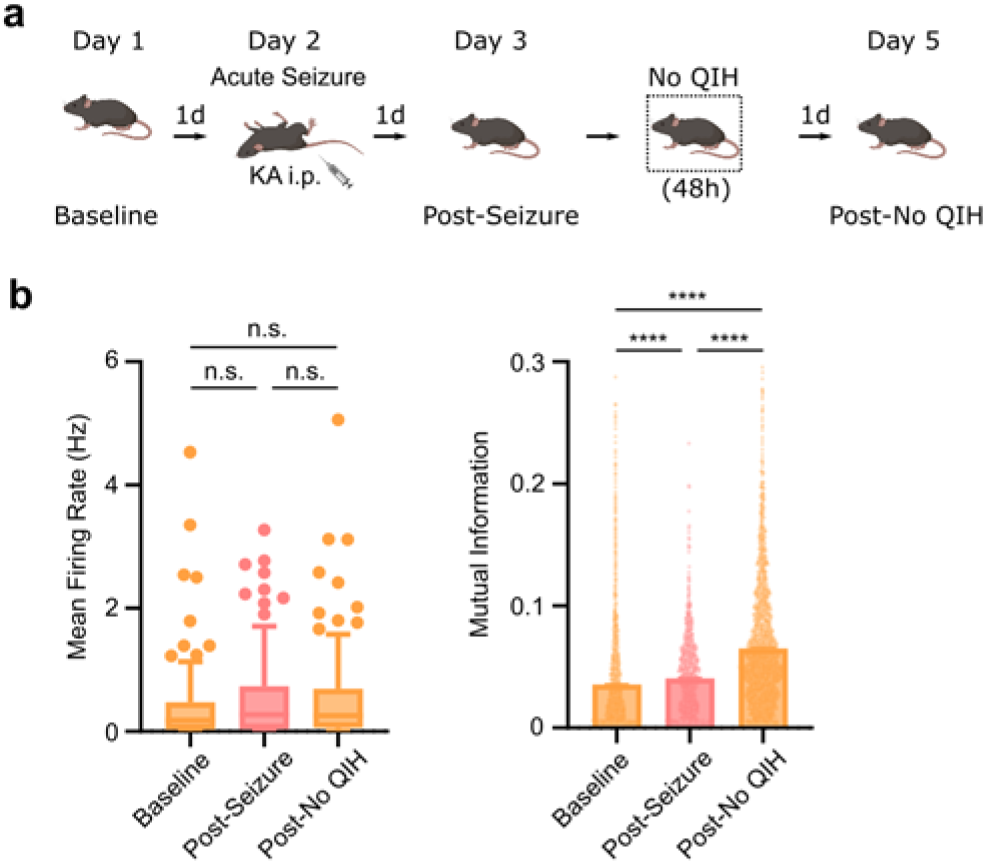
Firing rates and MI changes in the hippocampal CA1 neurons after seizure without QIH. **a,** Experimental design for the control experiment examining neuronal interaction changes following kainate-induced status epilepticus without subsequent QIH induction. CA1 activity was recorded at baseline, after KA-induced seizures, and after 48 hours of stay in the hibernation chamber without QIH induction (No QIH). **b,** Mean firing rate of individual units and pairwise MI at baseline, after seizures and after No QIH. KA-induced seizures increased pairwise MI without a corresponding persistent increase in mean firing rate (Kruskal-Wallis test, followed by post-hoc multiple comparisons with Dunn’s correction; non-matched units, n = 365 units from five mice; firing rate, p = 0.3070 for Baseline vs Post-Seizure, p = 0.2040 for Baseline vs Post-No QIH, and p > 0.9999 for Post-Seizure vs Post-No QIH; MI, p < 0.0001 for all comparisons).

**Extended Data Fig. 2.**
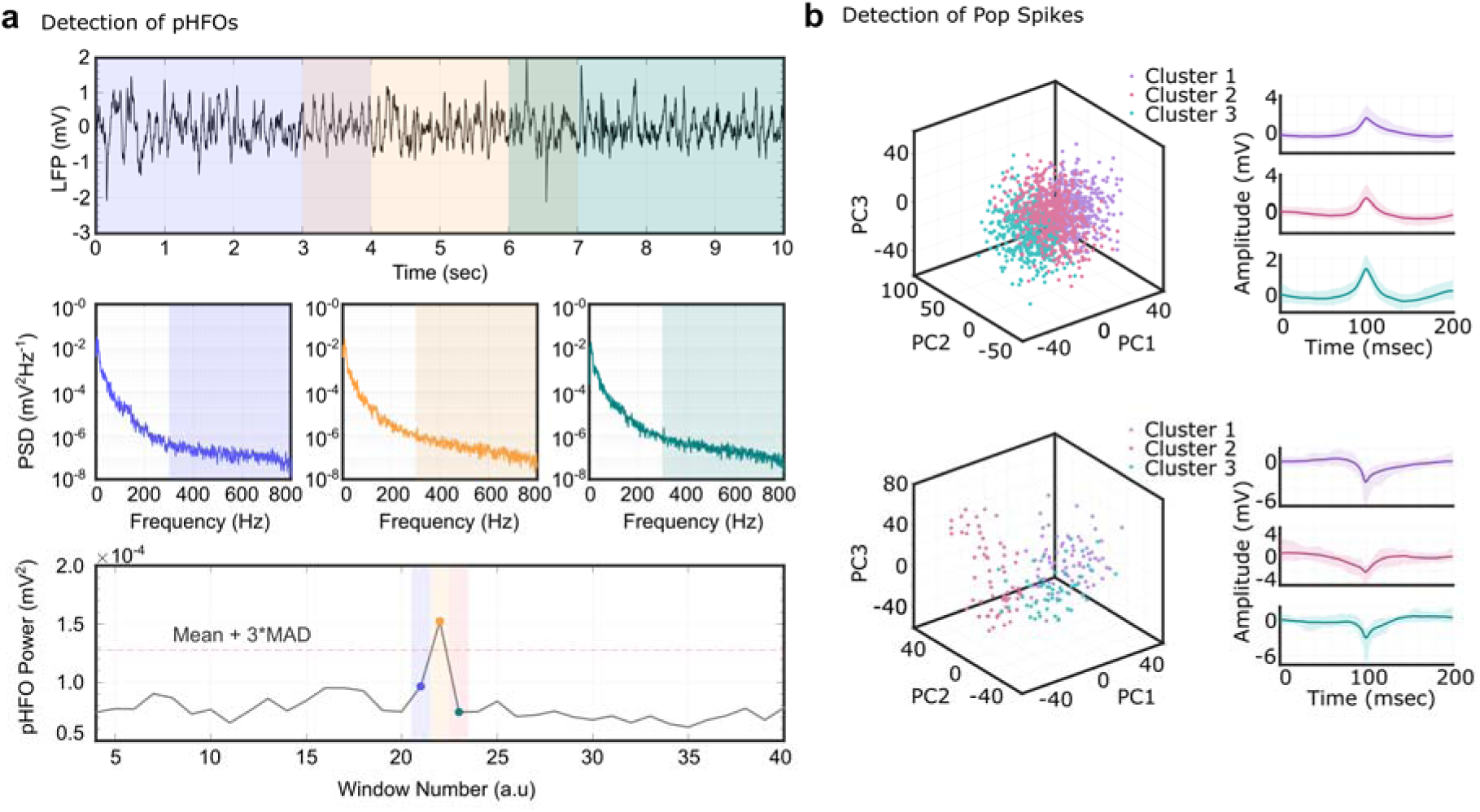
Detection of pathological LFP activities associated with the development of epilepsy. **a,** An illustration of the pHFO detection procedure using a 10-s LFP example segment (See Methods). The LFP was resampled at 1.6 kHz and browsed using overlapping 4-s windows with 1-s overlap (0∼4 s in shaded blue, 3∼7 s shaded orange, 6∼10 s shaded teal) (top). Power spectral density (PSD, Welch’s method) and the pHFO band (300∼800 Hz) are indicated by dashed lines (middle). Power in the pHFO band for 40 consecutive windows (bottom). Colored points correspond to the selected windows shown in the top and middle plots. A horizontal dashed red line marks the detection threshold (median + 3×MAD), above which windows were classified as HFO events.). **b,** For each automatically detected population spike, a 200-ms waveform was extracted and analyzed. Waveforms were grouped using principal component analysis (PCA) followed by k-means clustering (see Methods). (left) 3D scatter plot of the first three PCA components, with points colored according to cluster assignment. (right) Mean waveform (solid line) and 5–95% percentile envelope (shaded area) for each cluster. Through this approach, both positive and negative pop spikes were identified and pooled for the analysis in the main text.

**Extended Data Video. 1.**
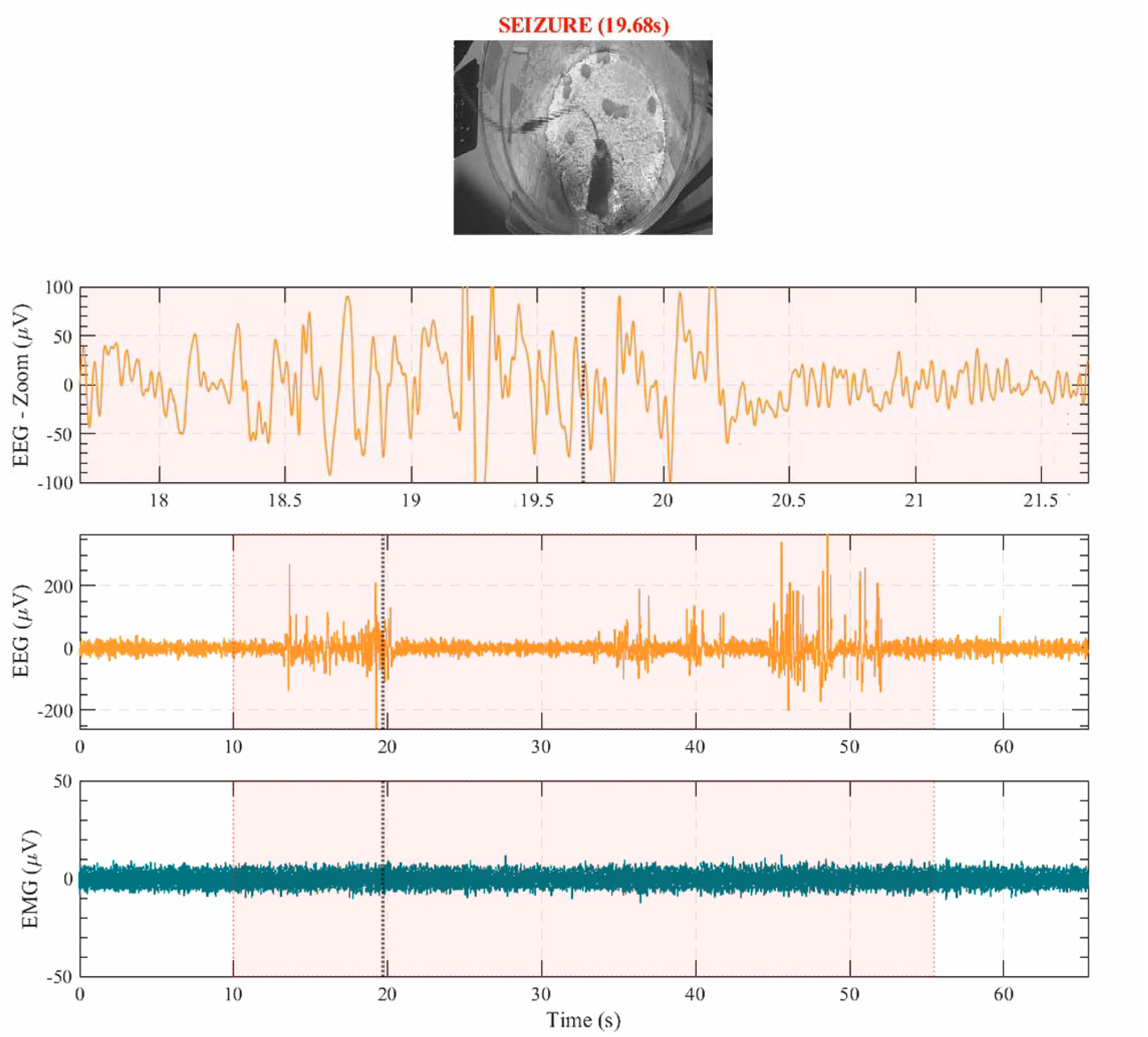
Representative recurrent seizure events (EEG, EMG, and behavior) Top panel: behavioral video synchronized to electrophysiological recordings. Lower panels: EEG (orange; average of two channels) and EMG (teal). EEG is shown as (i) a sliding 4 s window centered on the current time point and (ii) The full-scale trace. Vertical dashed lines indicate the current time point. Seizure epochs (EEG abnormalities without EMG activity increase) are marked by a semi-transparent red overlay. Noticeable seizures : 10s ∼ 20s : EEG power: moderate, behavior severity : low; 35s ∼ 42s : EEG power: moderate, behavior severity : low; 44s ∼ 52s : EEG power: high, behavior severity : high

