## Supplementary figures and images for "Artificial hibernation uncovers distinct interaction dynamics in functional and pathological state change"

### Extended Data Figure 1

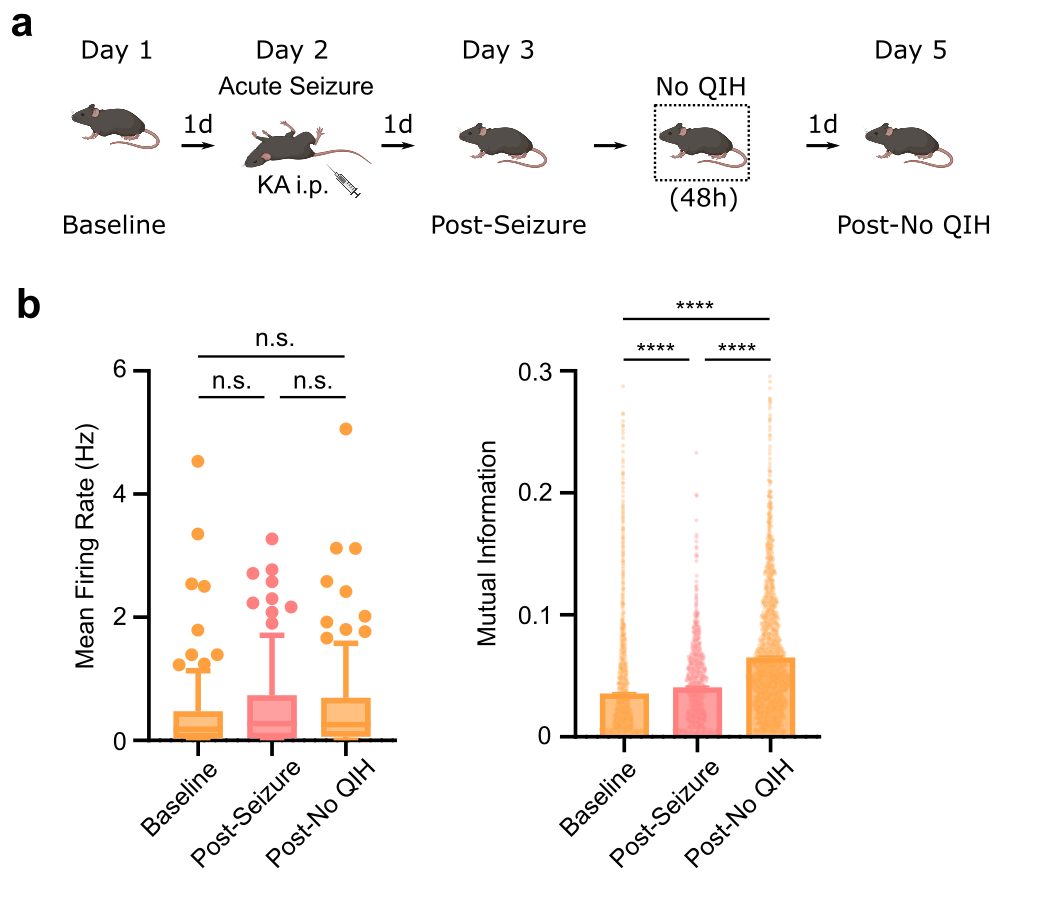

### Extended Data Figure 2

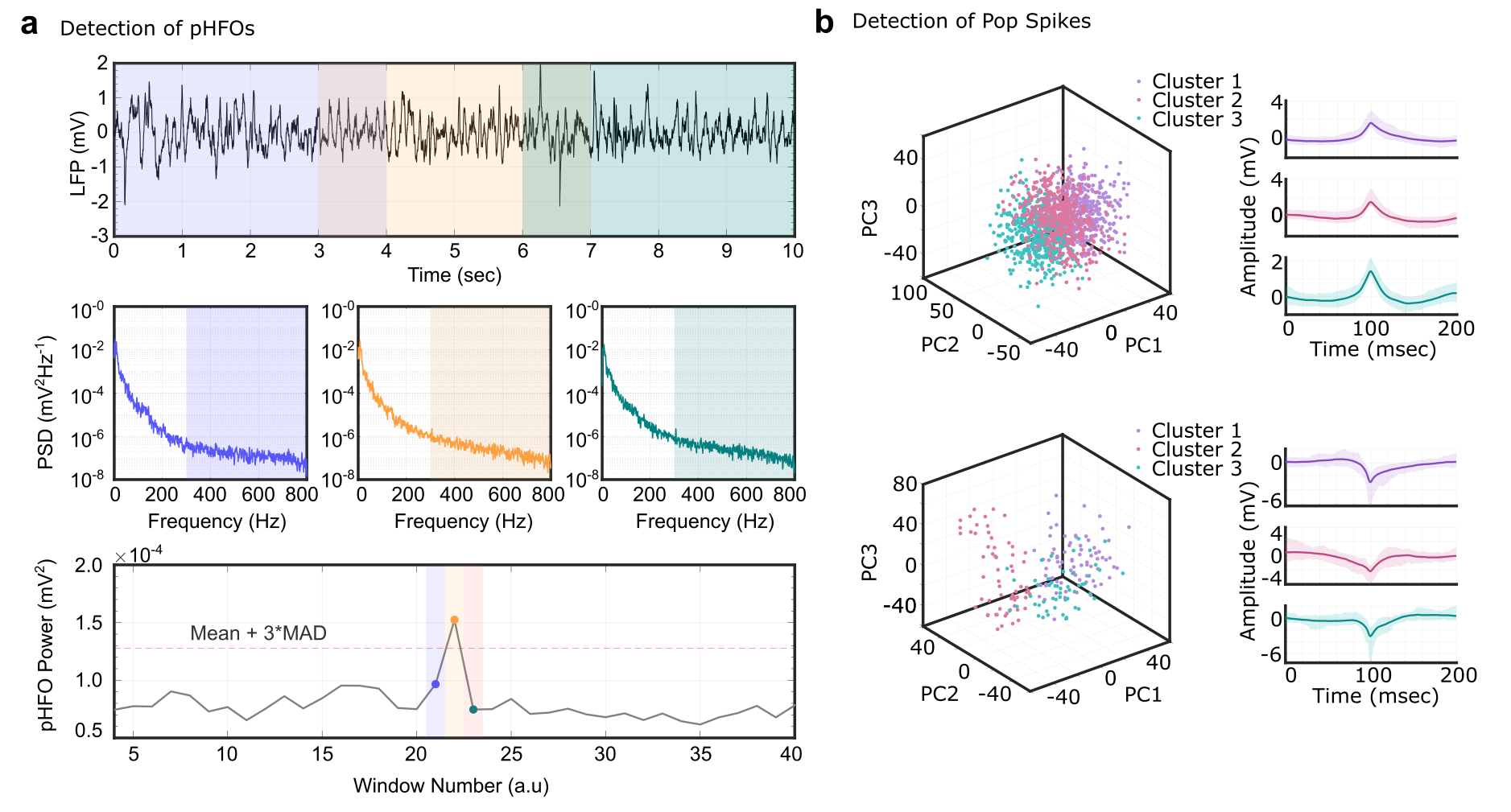
